# The effects of environmental heterogeneity within a city on the evolution of clines

**DOI:** 10.1101/2022.04.06.487365

**Authors:** James S. Santangelo, Cindy Roux, Marc T. J. Johnson

## Abstract

1. There is increasing evidence that environmental change associated with urbanization can drive rapid adaptation. However, most studies of urban adaptation have focused on coarse urban vs. rural comparisons or sampled along a single urban-rural environmental gradient, thereby ignoring the role that within-city environmental heterogeneity might play in adaptation to urban environments.
2. In this study, we examined fine-scale variation in the presence of HCN—a potent anti-herbivore defense—and its two underlying genes (*Ac* and *Li*) between park green spaces and surrounding suburban habitats for five city parks in the Greater Toronto Area.
3. We show that fine-scale urbanization has driven the formation of micro-clines in HCN on a scale of < 2 km, though the presence and strength of micro-clines varied across parks. Interestingly, these micro-clines were driven by lower HCN frequencies inside park green spaces, and are therefore in the opposite direction to that predicted based on previously described patterns of HCN frequency change along urban-rural gradients.
4. *Synthesis:* These results suggest larger scale, adaptive urban-rural clines occur across a complex matrix of environmental heterogeneity within cities that drives fine-scale adaptive microclines of varying strengths and directions.

## Introduction

Global urbanization is causing dramatic changes to the physical environment and the biota living within cities (Grimm *et al*. 2008). Recently, urban areas have been shown to be a potent driver of evolutionary change, capable of affecting all principle mechanisms of evolution, including mutation, genetic drift, gene flow, and natural selection (Johnson & Munshi-South 2017; Szulkin *et al*. 2020b). An increasing number of studies are treating global urbanization as a large-scale unplanned experiment, as a way to study parallel evolution (Perrier *et al*. 2020; Santangelo *et al*. 2020a; Cosentino & Gibbs 2022), and in particular adaptation to anthropogenically mediated environmental change (Mueller *et al*. 2013; Donihue & Lambert 2014; Alberti *et al*. 2016; Winchell *et al*. 2016). The majority of studies of adaptation to urbanization treat urban areas as either homogeneous environments, contrasting urban versus nonurban areas (Winchell *et al*. 2016; Theodorou *et al*. 2018; Szulkin *et al*. 2020a), or as continuous gradients of environmental change (Santangelo *et al*. 2020b). The reality is that the unique effects of human activity and behaviour on physical and biotic factors causes urban environments to be highly heterogeneous in space and time at the scale of a single city (Cadenasso *et al*. 2007; Pickett *et al*. 2017), and the evolutionary consequences of such heterogeneity for adaptation within cities is unknown (Rivkin *et al*. 2019; Alberti *et al*. 2020). Here we focus on the importance of spatial heterogeneity within cities, what we call intraurban heterogeneity, for phenotypic evolution.

Environmental heterogeneity within cities has the potential to lead to fine-scale local adaptation of populations. Intraurban heterogeneity comes in many forms, including variation in the amount and distribution of green space (e.g., parks), blue space (e.g., ponds, rivers, and shorelines), grey infrastructure (e.g., residential, industrial, roads and railways), biotic interactions, as well as social heterogeneity in human populations (e.g., density, economic, cultural and race) (Tanner *et al*. 2014; Pickett *et al*. 2017; Des Roches *et al*. 2020; Schell *et al*. 2020). It is well established that the heterogeneity across urban areas affects genetic drift and gene flow (Munshi-South 2012; Munshi-South *et al*. 2016; Lourenço *et al*. 2017; Beninde *et al*. 2018), influencing the amount of genetic diversity within populations and the movement of alleles and genetic differentiation between populations (Miles *et al*. 2019; Schmidt *et al*. 2020). But how such heterogeneity influences natural selection and ensuing adaptive evolution is poorly understood (Gorton *et al*. 2018; Rivkin *et al*. 2019). Classical theoretical models demonstrate how populations can adapt to their local environment, even in the face of gene flow (May *et al*. 1975; Slatkin 1975; Lenormand 2002). These theoretical predictions are well supported by numerous studies of local adaptation over small spatial scales in nonurban environments (Antonovics & Bradshaw 1970; Snaydon & Davies 1972; Hoekstra *et al*. 2006; Tack & Roslin 2010; Selby & Willis 2018). This body of theory and empirical work provides a framework to study how environmental heterogeneity caused by the unique anthropogenic disturbance of urban development may lead to fine-scale adaptation.

Here we test the hypothesis that intraurban environmental heterogeneity created by environmental differences between city parks and surrounding suburban areas drives repeated fine-scale phenotypic clines consistent with adaptation. We tested this hypothesis using the white clover system (*Trifolium repens*) because it has already been shown to exhibit repeated adaptive clines along urban-rural gradients across cities throughout the world in an ecologically important trait (hydrogen cyanide—HCN), as well as the Mendelian inherited loci underlying the trait (Santangelo *et al*. 2020b, 2022). HCN is a potent anti-herbivore defense that also affects tolerance to abiotic stressors. Specifically, current evidence suggests a benefit to producing HCN and/or its metabolic components under moderate to frequent drought stress (Kooyers & Olsen 2013; Kooyers *et al*. 2014), and a cost to their production in frost-prone environments (Kooyers *et al*. 2018). We may therefore expect higher HCN frequencies in habitats with greater herbivore damage, and higher summer and winter temperatures leading to greater drought stress and reduced frost exposure, respectively. We used a unique geological feature of the Greater Toronto Area, parallel ravines across the city, because they create naturally replicated parks that have similar transitions from green parks to surrounding dense residential areas. We further sought to identify the abiotic and biotic environmental drivers of genetic clines, including how % impervious surface, herbivory, and temperature are related to changes in HCN, and alleles at individual loci.

## Materials and Methods

### Study System

White clover (*Trifolium repens* L.) is a classic system for the study of adaptive clines to environmental gradients. *Trifolium repens* contains a Mendelian polymorphism for the production of the antiherbivore defence hydrogen cyanide (HCN), which is either present or completely absent in plants (Corkill 1942; Hughes 1991). This variation in HCN results from allelic variation at each of two loci: a closely linked three gene cluster (hereafter *Ac*) involved in the biosynthesis of cyanogenic glycosides (i.e., linamarin and lotaustralin), and *Li*, which encodes for linamarase (Olsen *et al*. 2007, 2008; Olsen & Small 2018). These components are stored in separate cellular compartments, and when the cell is damaged by herbivores or freezing, linamarase cleaves the glycoside to produce HCN. Both *Ac* and *Li* exhibit partial or whole gene deletions, creating the nonfunctional recessive alleles *ac* and *li*, respectively (Olsen *et al*. 2013). A plant requires at least one functional copy of both *Ac* and *Li* to produce HCN, and individuals that are homozygous for either *ac* or *li* are acyanogenic.

While HCN is a potent antiherbivore defence (Dirzo & Harper 1982; Thompson & Johnson 2016; Santangelo *et al*. 2018b), HCN and/or the presence of functional *Ac* and *Li* alleles decreases tolerance to freezing (Daday 1965; Brighton & Horne 1977; Olsen *et al*. 2008; Kooyers *et al*. 2018). Daday (Daday 1954b, a) showed decreased frequencies of HCN and functional *Ac* and *Li* alleles at higher latitudes and altitudes where temperatures are lower. These clines occur repeatedly in both the native and introduced range of *T. repens* (Daday 1958), which is interpreted as evidence for adaptive clines caused by spatial variation in selection for the benefits of defence, and against the costs associated with reduced freezing tolerance in cyanogenic plants (Hughes 1991). In addition, experimental data suggest a benefit to producing cyanogenic glucosides in the presence of drought stress (Kooyers *et al*. 2014), leading to higher HCN frequencies in more arid environments (Kooyers & Olsen 2013; Kooyers *et al*. 2014). More recently it has been shown that similar gradients in HCN, *Ac* and *Li* evolve in response to environmental variation caused by urbanization, with repeated urban-rural clines across cities (Thompson *et al*. 2016; Johnson *et al*. 2018; Santangelo *et al*. 2020b). A combination of experiments (Thompson *et al*. 2016), population genetic analysis (Johnson *et al*. 2018; Santangelo *et al*. 2022), and modeling (Santangelo *et al*. 2018a), strongly support the conclusion that these clines are caused by adaptation to environmental gradients along urban-rural transects.

To test how intraurban spatial environmental heterogeneity may affect adaptation, we studied changes in the frequency of HCN, *Ac* and *Li* across five park-suburban gradients across the Greater Toronto Area (GTA). The GTA is Canada’s most densely populated region, with ~6M residents over an area of >7000 km^2^. Our approach was facilitated by a series of river valleys that run perpendicular to the GTA’s Lake Ontario shoreline. Most of these valleys have been converted to city parks, with a mixture of mowed grass, fields, and forest. Dense suburbs often surround the river valleys, leading to a stark transition from large green spaces to suburbs. We selected five parks (from West to East: Erindale Park along the Credit River; Humber Park along the Humber River; High Park adjacent to Grenadier Pond, a former glacial river; Riverdale Park along the Don River; Rouge Park along the Rouge River) to study whether *T. repens* exhibits fine scale clines in HCN, *Ac* or *Li* from green space to suburbs using the methods described below.

### Sampling

In summer 2019, we sampled white clover stolons throughout each park’s central green space, and along eastern and western transects spanning the park’s bordering suburbs (Figure 1). We focused on environmental heterogeneity represented by the stark ecotones created by the transition from large green parks to dense surburban housing developments. We chose these environments because they represent some of the greatest extremes in environments within cities, and thus were predicted to be the most likely to show phenotypic differentiation over fine spatial scales. Specifically, we sampled 96 individuals within each park’s green space, trying to maximize the amount of park surface covered while ensuring at least 3 m between adjacent plants to avoid sampling the same genet. Then, beginning at the park’s eastern or western border with the surrounding suburbs, we collected an additional 96 plants per transect with approximately 20 meters between adjacent plants resulting in approximately 2 km long eastern and western transects. The only exception was High Park, which is bordered on the west by a water body and thus contained only park and eastern transect samples. Within park green spaces, plants were collected in large public lawns, whereas suburban collections occurred primarily within private lawns and public boulevards along sidewalks. In both cases, lawns were frequently mowed and dominated by grasses, legumes (e.g, *Medicago*), and asters (e.g., *Taraxacum*); there were no obvious differences in plant communities or vegetation cover across sampling sites. In total, we collected 1,034 plants across the five parks, each of which contained at least 4 intact leaves. All leaves from each plant were placed in 2 mL microcentrifuge tubes (one tube per plant) and stored at −80ºC before HCN, *Ac*, and *Li* phenotyping.

**Figure 1:**
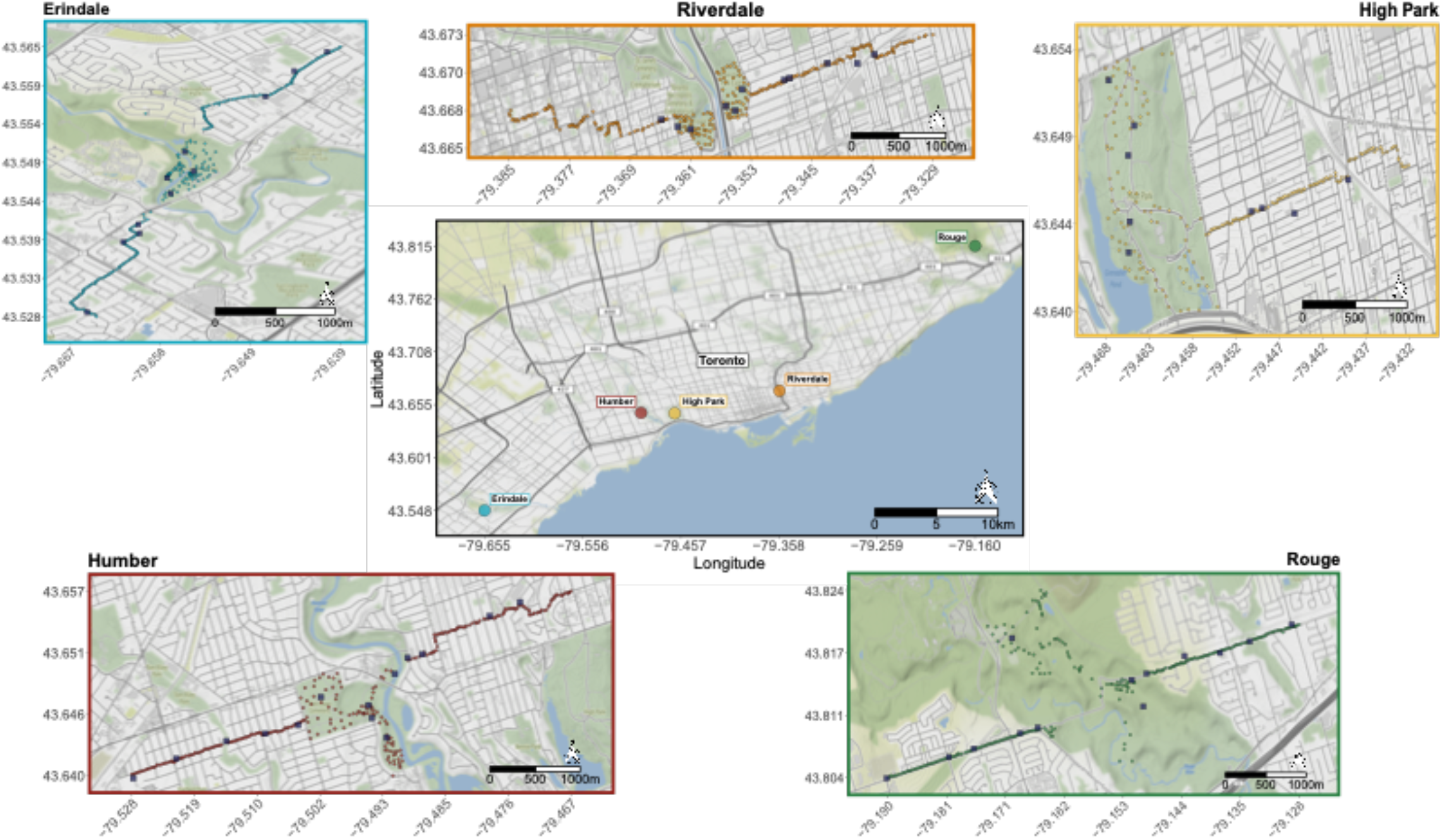
Map of sites sampled across the city of Toronto. Panels show the location of plants sampled within each site, and the iButtons purple squares) that were placed to record winter and summer temperatures within park green spaces and surrounding suburban transects.

### Herbivory assays

We quantified the amount of herbivore damage on each sampled plant to help unravel potential drivers of fine scale clines across urban parks. For each plant, we visually estimated the proportion leaf area eaten by herbivores on the third oldest, fully-expanded leaf to ensure enough time had elapsed to allow accumulation of herbivore damage and to provide leaf age-standardized estimates of herbivory across plants. Visual estimates of herbivory provide results consistent with those obtained from imaging software (Johnson *et al*. 2016) and are sufficiently precise to enable quantification of defense-meditated differences in herbivory in this system (Thompson & Johnson 2016; Santangelo *et al*. 2018b).

### Phenotyping

We used Feigl-Anger assays (Feigl & Anger 1966) with assay papers prepared according to Gleadow (*et al*. 2011) to determine the presence of HCN, *Ac*, and *Li* for each sampled plant. Briefly, we placed a single leaf from each sample in alternating wells in a 96-well PCR plate, added 80 μL of dH_2_0 to each well, macerated the tissue with a pipette tip to lyse the cells, placed the Feigl-Anger paper above the plate and incubated it for 3 hours at 37°C. The presence of a blue dot above the well indicates the presence of HCN for that sample and thus the presence of a dominant allele at both *Ac* and *Li* (genotype *Ac– Li–*). For plants that tested negative for HCN, we repeated the assay using either: (1) 80 μL of exogenous 0.2 EU/mL Linamarase (LGC Standards CDX-00012238-100) to test for the presence of cyanogenic glucosides (genotype *Ac*– *lili*) or (2) 50 μL dH_2_0 plus 20 μL of exogenous 10 mM Linamarin (Sigma-Aldrich 68264) to test for the presence of Linamarase (genotype *acac Li–*). Plants that are negative in all three tests lack dominant alleles at both loci (genotype *acac lili*).

### Temperature

Given evidence for the role of temperature in driving the formation of cyanogenesis clines on continental and local scales (Daday 1954a, 1965; Kooyers & Olsen 2013; Thompson *et al*. 2016; Santangelo *et al*. 2020b), we quantified ground temperature within parks and surrounding suburbs. Between the end of May and beginning of June 2019, we placed five iButton temperature probes (DS1922L iButtons, IButtonLink, Whitewater WI, USA) in shaded locations within each park and along each transect (N = 59), which recorded ambient temperature every 10 (Humber, High, Riverdale and Rouge Park), to 30 (Erindale) minutes. We only placed iButtons at a subset of the locations within park green spaces and surrounding suburban transects because it would have been infeasible to place them at every sampled plant, and our interest was to capture overall environmental differences between park green spaces and surrounding suburban transects. iButtons remained in the field continuously until late March 2020, except for a few days every 3 months to retrieve data and reset the probes. Apart from Erindale, we have detailed ambient temperature data for summer, fall, and winter for each park and surrounding transects. Due to a technical issue during the setup of the Erindale iButtons, we only have temperature data for part of the summer (May to July 2019) for this park and its associated urban transects. Of the 59 iButtons placed in the field, 32 remained during the first round of data collection and probe reset and 17 remained at the end of the experiment; the remaining iButtons were lost to demonic intrusion (e.g., vandalism and disappearance, Hurlbert 1984).

### Impervious surface

To better quantify the urban landscape surrounding sampled plants, we manually quantified the percentage of impervious surface around each plant. Using Google Earth Pro, we traced a 10-meter radius circle around each plant and within this circle, manually traced along the perimeter of all man-made impervious structures using the polygon tool. We then estimated the area of all outlined impervious surfaces and divided this by the total area of the circle (314 m^2^) to estimate the % impervious surface around each plant.

### Statistical analyses

To examine whether urban heterogeneity was associated with variation in fine-scale cyanogenesis clines, we first examined how the presence of HCN, *Ac*, or *Li* changes from park green spaces to surrounding urban transects, and whether this effect is consistent across different parks (hereafter “site”). Specifically, we began by fitting a binomial regression using the presence/absence of HCN, *Ac*, or *Li* as a response variable, and site (i.e., one of five sampled parks), habitat (i.e., park green space or suburban transect), and their interaction as predictors.

We then followed up on this model to better understand the environmental drivers of fine-scale variation in HCN and allele frequencies across sites. We again fit binomial regressions using the presence/absence of HCN, *Ac*, or *Li* as a response variable but instead used the following predictors: site, % impervious surface, herbivory, the site ⨉ impervious surface interaction, and the site ⨉ herbivory interaction. Model fitting and *P*-value estimation were performed as above. We additionally ran similar models to those above but using the proportion of herbivore damage as the response variable to examine how herbivory differed between green spaces and surrounding urban habitats. Finally, we ran models using distance as a predictor instead of % impervious surface (with distance set to zero for plants within park green spaces) for a more spatially informed model: these models yielded qualitatively identical conclusions, likely because distance and % impervious surface are correlated (*r*_*Pearson*_ = 0.70), and thus we present results with distance in the online supplementary materials (Text S1, TableS1). All binomial regressions were fitted using the *glm()* function in R v.4.0.3 (R Core Team 2020). *P*-values were obtained by fitting models with type III SS and re-fitting with type II SS in the absence of significant interaction (Langsrud 2003; Landsheer & van den Wittenboer 2015) using the *Anova()* function from the *car* package (Fox & Weisberg 2011). All diagnostics of model fits and their residuals were performed using the DHARMa package (Hartig 2022).

Given evidence that temperature is frequently associated with the frequency of HCN and underlying genes, we were interested in examining whether temperature differences between green spaces and surrounding urban transects could explain differentiation in HCN, *Ac*, or *Li* frequencies over those same spatial scales. We began by using daily iButton temperature readings to identify the coldest and the warmest months, reasoning that if temperature was an important selective agent in driving clines, its effects would be most apparent at its most extreme values. Using readings from all 59 iButtons, we defined the coldest month as the month with the lowest mean daily minimum temperature, and the warmest month as the month with the highest mean daily maximum temperature. From these criteria, we identified February as the coldest month and July as the warmest month. We then filtered iButton temperature readings to include only values in these months and fit mixed-effects models with mean daily minimum or maximum temperature as a response variable and site, habitat, and their interaction as fixed-effects, and random-intercepts for each iButton. We also ran models replacing habitat with % impervious surface. We then qualitatively assessed whether differences in temperature could explain differences in HCN or gene frequencies between parks and surrounding urban transects. Model fitting and diagnostics were performed as above.

## Results

### Presence of Clines

The frequency of HCN and *Ac*, but not *Li*, differed between green spaces and surrounding suburban transects, but this effect varied among sites. Across all sites, the probability that a plant produced HCN increased by 50% from the lowest % impervious surface [0%, inside park green space, P(HCN+) = 0.23] to the highest % impervious surface [96.5%, P(HCN+) = 0.34] (main effect of % impervious surface: *X*^2^_1_ = 5.48, *P* = 0.02, Fig. 2D, Fig. S1, Table 1). Similarly, *Ac* increased by 37% across our gradient of impervious surface (main effect of % impervious surface: *X*^2^_1_ = 12.8, *P* < 0.001, Fig. 2E, Fig. S2, Table 1). However, the effects of green spaces (Site ⨉ Habitat interaction; HCN: *X*^2^_4_ = 10.5, *P* = 0.03; *Ac*: *X*^2^_4_ = 32.08, *P* < 0.001, Fig. 2A, B, Table 1), and impervious surface (Site ⨉ Impervious surface interaction: HCN: *X*^2^_4_ = 8.39, *P* = 0.08; *Ac*: *X*^2^_4_ = 22.49, *P* < 0.001, Fig 2D, E, Table 1), varied considerably among parks, and were largely driven by strong clines in High Park and Riverdale. Indeed, apart from Erindale, all sites tended to have less HCN along suburban transects. However, we avoid interpreting this difference since the error bars are largely overlapping (Fig. 2). By contrast, *Li* frequencies did not differ between green spaces and suburban areas (Fig 2C, Table 1) or with impervious surface gradients in any site (Fig. 2E, Fig. S3, Table 1).

**Figure 2:**
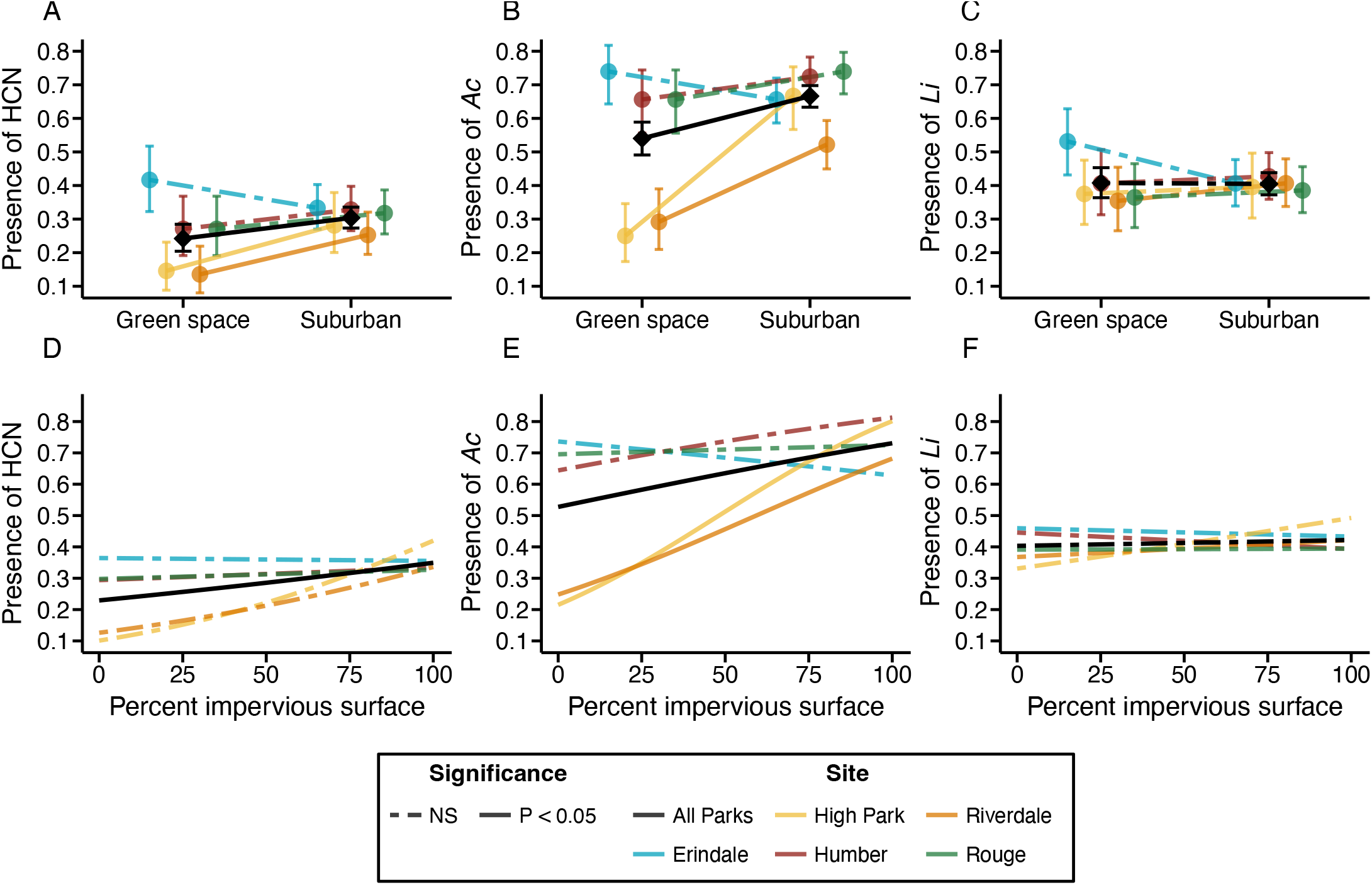
Presence of HCN, *Ac*, and *Li* as a function of habitat (i.e., Park green space vs. urban transects, A - C) or % impervious surface (D F). Colored lines show reaction norms (A - C) or clines (D - F) for each of the five sampled sites. Black diamonds (A - C) and solid black nes show the main effect of habitat or % impervious surface across all sites. Where significant interactions were detected, solid lines denote dividual site reaction norms or clines that are significant at *P* < 0.05 following *post-hoc* contrasts.

**Table 1:** Effects of intraurban heterogeneity on HCN defences and the presence of *Ac* and *Li* alleles. We report *X*_*2*_ statistics, degrees-of-freedom, and *P-*values for independent variables in models predicting the presence/absence of HCN, *Ac*, or *Li*. Models were run treating habitat as binary (i.e., park green space vs. suburbs) or using per-plant estimates of % impervious surface and herbivory. Bolded terms are significant at *P* < 0.05. “Site” refers to the five parks sampled across the city. % impervious surface and herbivory were estimated for each sampled plant.

| Response | Predictor | $X^2$ | df | $P$ |
| --- | --- | --- | --- | --- |
| *Models with green space vs. suburbs* |  |  |  |  |
| HCN | Site | **27.4** | **4** | **< 0.001** |
|  | Habitat | **6.59** | **1** | **0.01** |
|  | Site × Habitat | **10.51** | **4** | **0.03** |
| Ac | Site | **83.86** | **1** | **< 0.001** |
|  | Habitat | **23.8** | **1** | **< 0.001** |
|  | Site × Habitat | **32.08** | **4** | **< 0.001** |
| Li | Site | 3.9 | 4 | 0.41 |
|  | Habitat | 0.01 | 1 | 0.9 |
|  | Site × Habitat | 5.07 | 4 | 0.28 |
| *Models with impervious surface and herbivory* |  |  |  |  |
| HCN | Site | **21.79** | **4** | **< 0.001** |
|  | % Impervious | **5.48** | **1** | **0.02** |
|  | Herbivory | 0.02 | 1 | 0.88 |
|  | Site × % Impervious | 8.39 | 4 | 0.07 |
|  | Site × Herbivory | 1.23 | 4 | 0.87 |
|  | % Impervious × Herbivory | 2.2 | 1 | 0.14 |
|  | Site × % Impervious × Herbivory | 2.95 | 4 | 0.57 |
| Ac | Site | **64.4** | **4** | **< 0.001** |
|  | % Impervious | **12.8** | **1** | **< 0.001** |
|  | Herbivory | 0.14 | 1 | 0.71 |
|  | Site × % Impervious | **22.49** | **4** | **< 0.001** |
|  | Site × Herbivory | 0.44 | 4 | 0.98 |
|  | % Impervious × Herbivory | 1.19 | 1 | 0.27 |
|  | Site × % Impervious × Herbivory | 0.91 | 4 | 0.92 |
| Li | Site | 7.53 | 4 | 0.11 |
|  | % Impervious | 0.05 | 1 | 0.82 |
|  | Herbivory | 0.13 | 1 | 0.72 |
|  | Site × % Impervious | 5.14 | 4 | 0.27 |
|  | Site × Herbivory | **9.82** | **4** | **0.04** |
|  | % Impervious × Herbivory | 0.24 | 1 | 0.62 |
|  | Site × % Impervious × Herbivory | 4.7 | 4 | 0.32 |

### Effects of city parks on herbivory

Across all sites, herbivore damage was greater in park green spaces than in surrounding suburban habitats (main effect of habitat: *X*^2^_4_ = 4.11, *P* < 0.043, Fig. 3A, Table 2) and decreased by 67% across our gradient of impervious surface (main effect of impervious surface: *X*^2^_4_ = 4.06, *P* < 0.044, Table 2). This pattern of increased herbivore damage in park green spaces was consistent across sites as seen by the non-significant site ⨉ habitat and site ⨉ impervious surface interactions (Table 2). Herbivory did not predict variation in HCN, *Ac*, or *Li* frequencies in any site (all *P* > 0.05).

**Table 2:** Effects of intraurban heterogeneity on herbivory and summer/winter temperatures. We report *X*_2_ statistics, degrees of freedom, and *P-*values for independent variables in models predicting the herbivory, maximum summer temperature, or minimum winter temperature. Models were run treating habitat as binary (i.e., park green space vs. suburbs) or using per-plant estimates of % impervious surface and herbivory. Bolded terms are significant at *P* < 0.05. “Site” refers to the five parks sampled across the city. % impervious surface and was estimated for each sampled plant.

| Response | Predictor | $X^2$ | Df | $P$ |
| --- | --- | --- | --- | --- |
| *Models with green space vs. suburbs* |  |  |  |  |
| **Herbivory** | **Site** | **10.29** | **4** | **0.04** |
|  | **Habitat** | **4.11** | **1** | **0.04** |
|  | Site × Habitat | 0.98 | 4 | 0.91 |
| **Summer Max Temp** | Site | 8.99 | 4 | 0.06 |
|  | **Habitat** | **3.86** | **1** | **0.05** |
|  | Site × Habitat | 8.21 | 4 | 0.08 |
| **Winter Min Temp** | Site | 1.86 | 2 | 0.39 |
|  | Habitat | 1.39 | 1 | 0.23 |
|  | Site × Habitat | 1.5 | 2 | 0.47 |
| *Models with impervious surface and herbivory* |  |  |  |  |
| **Herbivory** | Site | 8.84 | 4 | 0.07 |
|  | **% Imperv** | **4.06** | **1** | **0.04** |
|  | Site × % Imperv | 0.65 | 4 | 0.96 |
| **Summer Max Temp** | Site | 5.5 | 4 | 0.23 |
|  | **% Imperv** | **7.43** | **1** | **0.006** |
|  | **Site × % Imperv** | **13.25** | **4** | **0.01** |
| **Winter Min Temp** | Site | 2.85 | 2 | 0.24 |
|  | % Imperv | 1.5 | 1 | 0.22 |
|  | Site × % Imperv | 1.44 | 2 | 0.48 |

**Figure 3:**
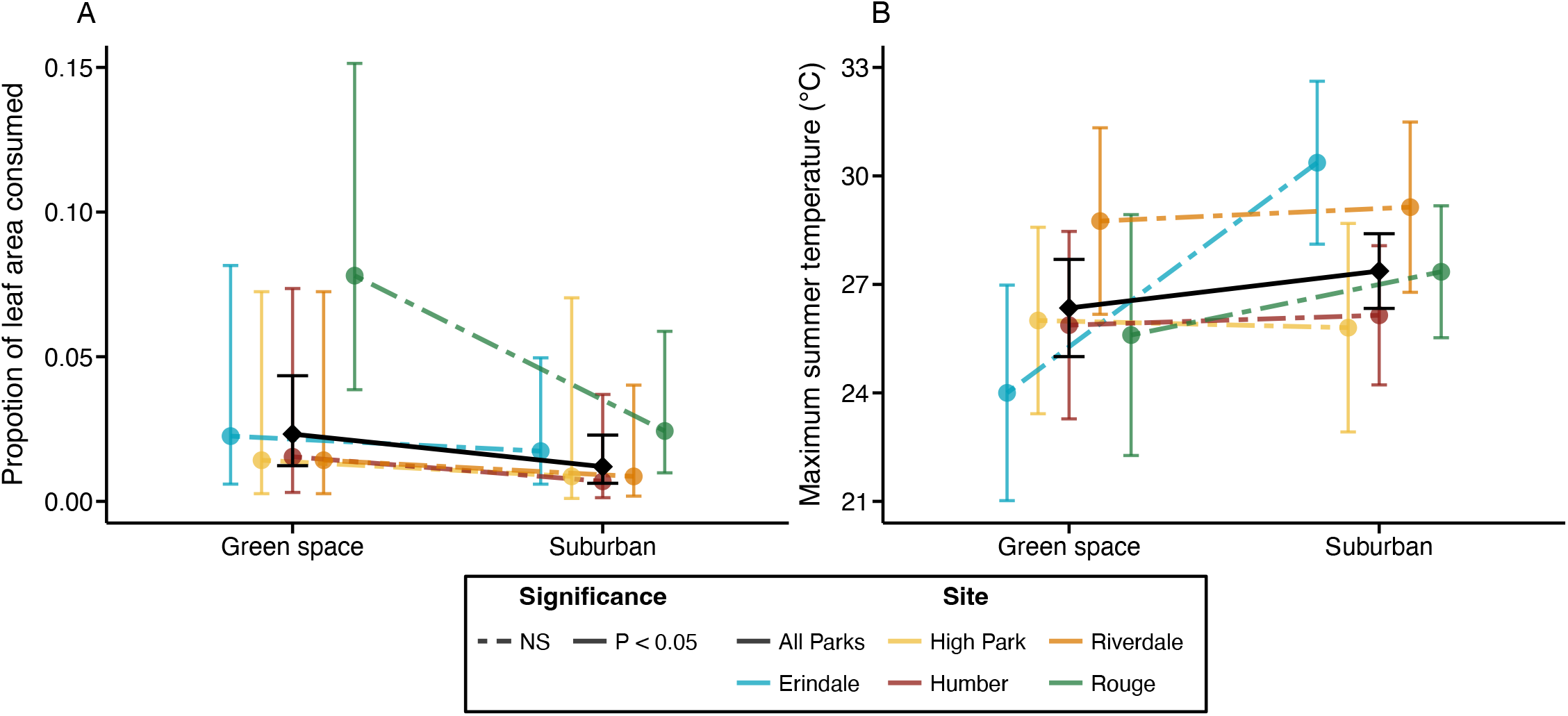
Reaction norms showing the change in herbivory (A) or maximum summer temperature (B) between green spaces and urban ansects for each of the five sites.

### Effects of city parks on temperature

Maximum summer temperature was on average lower in populations with lower impervious surface, but this effect varied across sites. Specifically, maximum summer temperature increased by 55% and 20% across our gradient of impervious surface in Erindale and Rouge Park, respectively, with little change in the other three sites (Site ⨉ Impervious surface interaction: *X*^2^_4_ = 13.30, *P* < 0.01, Table 2, Fig. 3B). By contrast, neither green space nor % impervious surface explained variation in minimum winter temperature across any park (Table 2).

## Discussion

Our results show that environmental heterogeneity within a city can drive evolution that is consistent with adaptation over small spatial scales. We had previously shown that *T. repens* repeatedly exhibits adaptive clines in HCN and its underlying alleles across urban-rural gradients at the scale of entire cities at a global scale (Santangelo *et al*. 2020b, 2022). Here we show that such clines can evolve even within a city across gradients of impervious surface at a scale of < 2 km. These clines are heterogenous in their occurrence and magnitude, but when they occur they involve large shifts in the frequency of plants producing HCN and the dominant allele *Ac*. While we cannot definitively rule-out non-adaptive mechanisms (i.e., genetic drift and gene flow) as drivers of these clines, a couple of observations suggest they are likely adaptive. First, drift would be expected to drive clines in random directions, especially for alleles frequencies at individual loci (i.e., *Ac* and *Li*, (Santangelo *et al*. 2018a). Therefore, the fact that four of the five sites have clines in HCN, *Ac*, and *Li* trending in the same direction suggests they are driven by natural selection. Second, all current population genomic evidence suggests high genome-wide neutral diversity in urban white clover populations, and high levels of admixture and low differentiation among populations (Johnson *et al*. 2018; Santangelo *et al*. 2022). While these results were obtained in the context of larger-scale urban-rural gradients, they likely apply to the smaller spatial-scale examined here where white clover is incredibly abundant, and outcrossing is achieved by pollinators capable of moving across much of our sampled transects. Together, these results suggest that natural selection is likely driving the formation of fine-scale spatial clines in HCN. What is most surprising is that the direction of these microclines due to intraurban heterogeneity (i.e., increasing HCN and *Ac* frequency with increased impervious surface) was in the opposite direction than expected based on the larger scale urban-rural clines previously observed in Toronto (Thompson *et al*. 2016) and many other cities (Santangelo *et al*. 2020b, 2022), where we frequently observed a decrease in the frequency of HCN, *Ac* and *Li* with increasing urbanization. This does not refute our earlier findings of adaptation to urban-rural gradients, but instead shows that this larger scale adaptive cline occurs across a complex matrix of heterogeneity within cities that drives microclines that in some instances are opposite in direction to the larger scale patterns.

Interestingly, we only observed clines in the *Ac* locus and not the *Li* locus, with a corresponding weak cline in the HCN phenotype. The presence of clines at only one locus suggests that in the case of intraurban heterogeneity, *Ac* may frequently be the target of selection rather than the HCN phenotype itself. While we had previously observed clines in both loci along urban-rural gradients in Toronto (see Fig. S3 in Thompson *et al*. 2016), subsequent work across 16 cities in eastern North America showed that clines in *Ac* are twice as common as clines in *Li* (Santangelo *et al*. 2020b). This implies that while HCN production might be the target of selection in some instances, *Ac* may include alternative biological functions that are targeted by selection in other cases. This raises the obvious question: What environmental factors might vary over fine spatial scales to select on these hypothesized alternative functionsã

A goal of our study was to identify the potential environmental drivers of microclines in HCN and its underlying alleles. While we detected changes in herbivore pressure and summer temperature between some green parks and surrounding suburbs, the direction of these changes, and the locations where they occurred, were inconsistent with the presence and direction of clines in HCN, *Ac* and *Li* (Table 1 and Table 2). Moreover, there was no clear difference in minimum winter temperature, an environmental stressor known to impose selection on HCN or its underlying alleles (Daday 1965; Thompson *et al*. 2016; Kooyers *et al*. 2018), including in Toronto (Thompson *et al*. 2016). Thus, it seems unlikely that differences in herbivory or temperature explain the microclines we observed. The higher frequency of the *Ac* allele in some suburbs is consistent with the hypothesized function of cyanogenic glycosides as nutrient storage molecules, especially under drought conditions (Kooyers *et al*. 2014). Suburban development frequently leads to decreased soil water content, lower soil nutrients and increased plant water stress (Marcotullio *et al*. 2008), which is expected to select for higher *Ac* frequency (Kooyers *et al*. 2014). Furthermore, a recent global study of urban adaptation showed that in the rare cases when HCN evolved higher levels in cities, those cities had greater potential for plant water stress in urban than rural areas, also likely due to selection to produce cyanogenic glucosides (Santangelo *et al*. 2022). Future work should seek to directly measure plant water stress and soil moisture in the field rather than relying on proxies such as temperature or satellite-derived estimates of aridity to better capture the role of this putative selective agent in driving cyanogenesis clines within cities and across urban-rural gradients.

Our study shows that intraurban environmental heterogeneity can drive fine-scale adaptive clines within cities. Moreover, the direction of these clines can in some instances seemingly oppose the direction of larger scale adaptive dynamics that occur at the scale of entire cities (Thompson *et al*. 2016) or even continents (Daday 1958). While it is true that urbanization is on average leading to increased homogenization of biotic and abiotic environmental factors at a global scale (McKinney 2006; Groffman *et al*. 2014; Santangelo *et al*. 2022), cities still contain stark environmental heterogeneity within them (Cadenasso *et al*. 2007; Pickett *et al*. 2017). This is most apparent in the stark transition from green parks to the surrounding suburban and urban matrix. Such dramatic environmental transitions are akin to the transitions that occur due to soil pollution created by mine tailings (Antonovics & Bradshaw 1970), fertilization and mowing regimes associated with agriculture (Snaydon & Davies 1972; Law *et al*. 1977), or natural changes in soil chemistry (Selby & Willis 2018). There are many additional types of environmental heterogeneity within cities (e.g., age of development, housing density, mowing and irrigation frequency, etc.), that vary at a diversity of spatial scales. Given that we see evidence of phenotypic differentiation along park-suburban ecotones, we propose it is likely that these other forms of complex heterogeneity are also likely to impose divergent selection. Whether or not urban populations will frequently adapt to intraurban environmental heterogeneity will depend on whether the predicted adaptive response of populations (i.e., based on the strength of selection and amount of standing genetic variation) can overcome the homogenizing effects of gene flow over such small spatial scales, or the stochastic changes in allele frequencies caused by genetic drift that are especially likely for small urban populations.

## Acknowledgements

We thank B. Cohan and S. Munim for help in conducting fieldwork.

