## Supplementary material for "The effects of environmental heterogeneity within a city on the evolution of clines": Fig. S

### Supplemental materials for:
### The effects of environmental heterogeneity within a city on the evolution of clines

**Contents:**


- Supplementary text S1
- Supplementary table S1
- Supplemental figures S1 to S3

**Supplementary text**

Text S1: Models with distance instead of % impervious surface

We ran the same models as those detailed in the main text but replacing % impervious surface with distance from park boundary for a mode spatially informed model. In these models, the distance of plants within park green spaces was set to 0, while the distance of plants along suburban transects was estimated as their distance from the nearest park edge using the Haversine distance.

The frequency of HCN and *Ac*, but not *Li*, differed between green spaces and surrounding suburban transects, but this effect varied among sites. Across all sites, the probability that a plant produced HCN increased from within park green spaces to surrounding suburban transects (main effect of distance: $X^2_1 = 4.97$, $P = 0.03$, Table S1). Similarly, *Ac* increased from within park green spaces to surrounding suburban transects (main effect of distance: $X^2_1 = 16.40$, $P < 0.001$, Table S1). However, the effects of distance (Distance × Site: HCN: $X^2_4 = 9.62$, $P = 0.47$; *Ac*: $X^2_4 = 19.90$, $P < 0.001$, Table S1), varied considerably among parks, and were largely driven by strong clines in High Park and Riverdale. By contrast, distance had no effect on *Li* frequencies at any sampled site (Table S1).

#### Supplemental tables

**Table 1:** Effects of intraurban heterogeneity on HCN defences and the presence of *Ac* and *Li* alleles. We report $X^2$ statistics, degrees-of-freedom, and *P*-values for independent variables in models predicting the presence/absence of HCN, *Ac,* or *Li*. Models were run with herbivory and distance as predictors, with the distance of plants within park green spaces set to zero. Bolded terms are significant at $P < 0.05$. "Site" refers to the five parks sampled across the city.

| Response | Predictor | $X^2$ | df | *P* |
|---|---|---|---|---|
| **HCN** | **Site** | **23.80** | **4** | **< 0.001** |
| | **Distance** | **4.97** | **1** | **0.03** |
| | Herbivory | 0.13 | 1 | 0.72 |
| | **Site × Distance** | **9.62** | 4 | **0.47** |
| | Site × Herbivory | 5.22 | 4 | 0.27 |
| | Distance × Herbivory | 0.15 | 1 | 0.70 |
| | **Site × Distance × Herbivory** | **10.36** | 4 | **0.03** |
| **Ac** | **Site** | **59.23** | **4** | **< 0.001** |
| | **Distance** | **16.40** | **1** | **< 0.001** |
| | Herbivory | 2.09 | 1 | 0.15 |
| | **Site × Distance** | **19.90** | **4** | **< 0.001** |
| | Site × Herbivory | 1.59 | 4 | 0.81 |
| | Distance × Herbivory | 0.52 | 1 | 0.47 |
| | Site × Distance × Herbivory | 4.90 | 4 | 0.30 |
| **Li** | Site | 3.77 | 4 | 0.48 |
| | Distance | 0.01 | 1 | 0.94 |
| | Herbivory | 0.01 | 1 | 0.91 |
| | Site × Distance | 0.97 | 4 | 0.92 |
| | **Site × Herbivory** | 8.89 | **4** | 0.21 |
| | Distance × Herbivory | 0.61 | 1 | 0.43 |
| | Site × Distance × Herbivory | 3.97 | 4 | 0.41 |

**Supplemental figures**

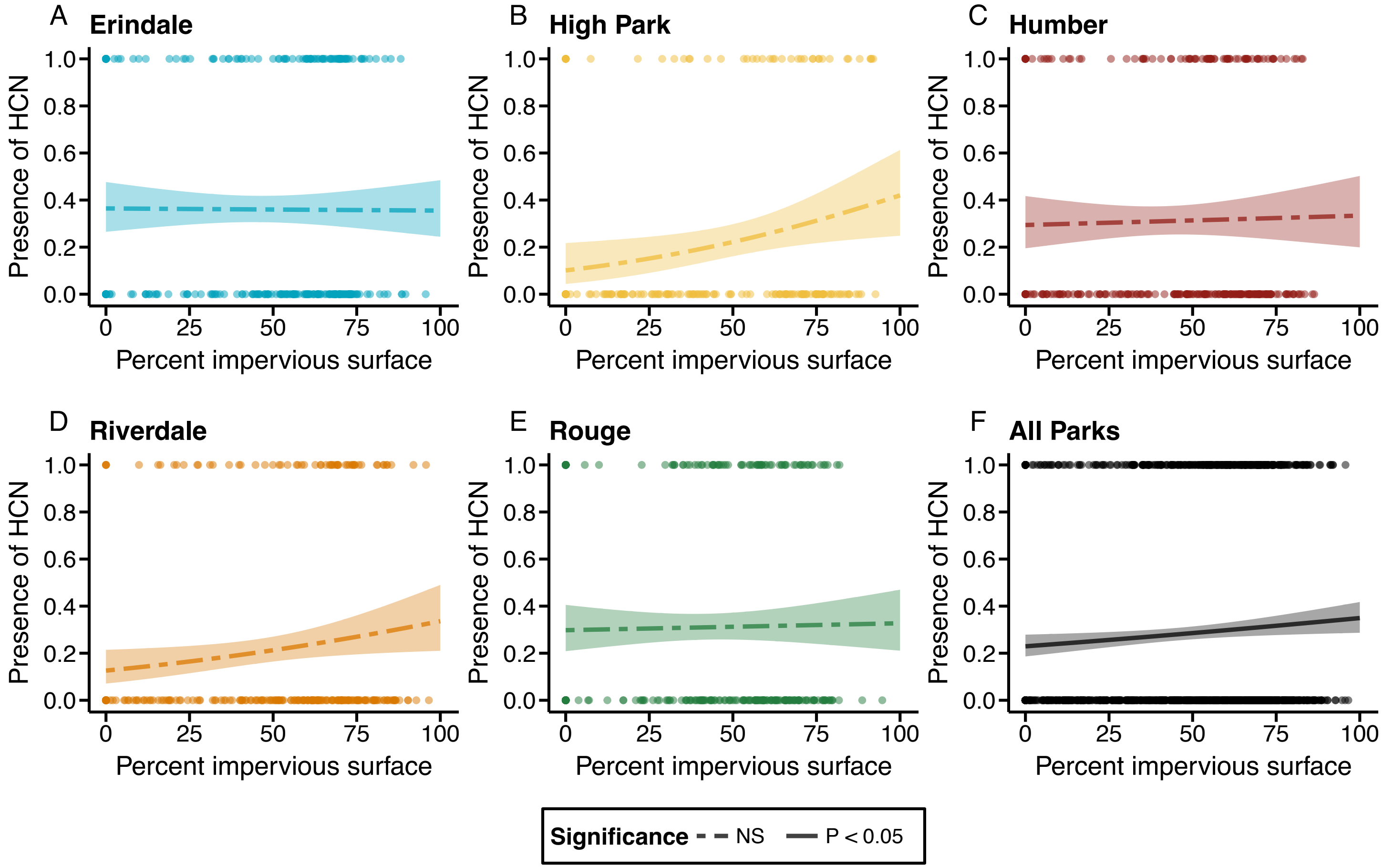


**Figure S1:** Clines in the frequency of HCN for each of the sampled sites (A – E) and across all sites combined (F). Solid lines represent significant clines ($P < 0.05$) based on a logistic regression of HCN presence/absence against percent impervious surface. Points represent individual plants. Shaded regions around lines represent the 95% confidence intervals around the fitted models.

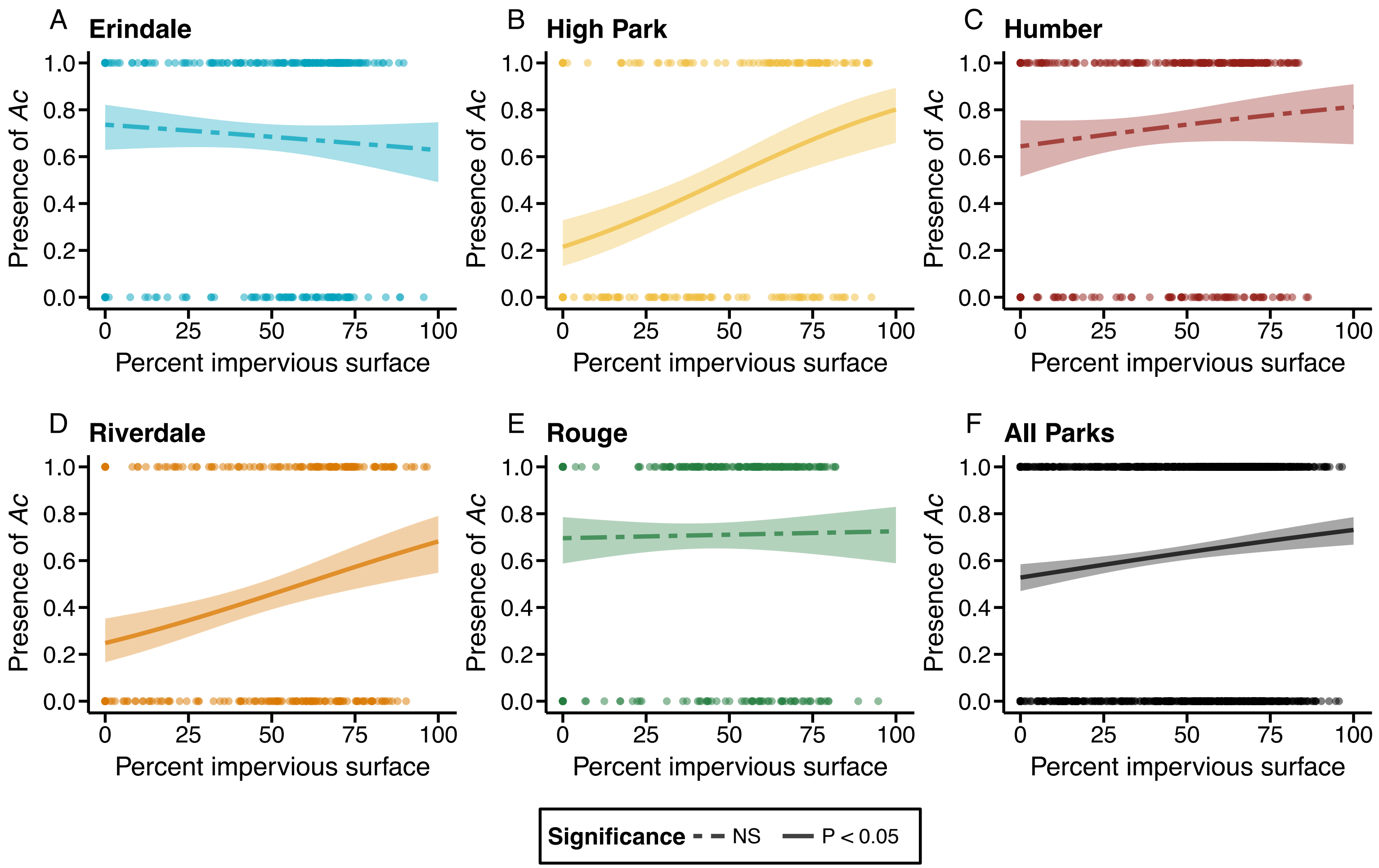


**Figure S2:** Clines in the frequency of *Ac* for each of the sampled sites (A – E) and across all sites combined (F). Solid lines represent significant clines ($P < 0.05$) based on a logistic regression of *Ac* presence/absence against percent impervious surface. Points represent individual plants. Shaded regions around lines represent the 95% confidence intervals around the fitted models.

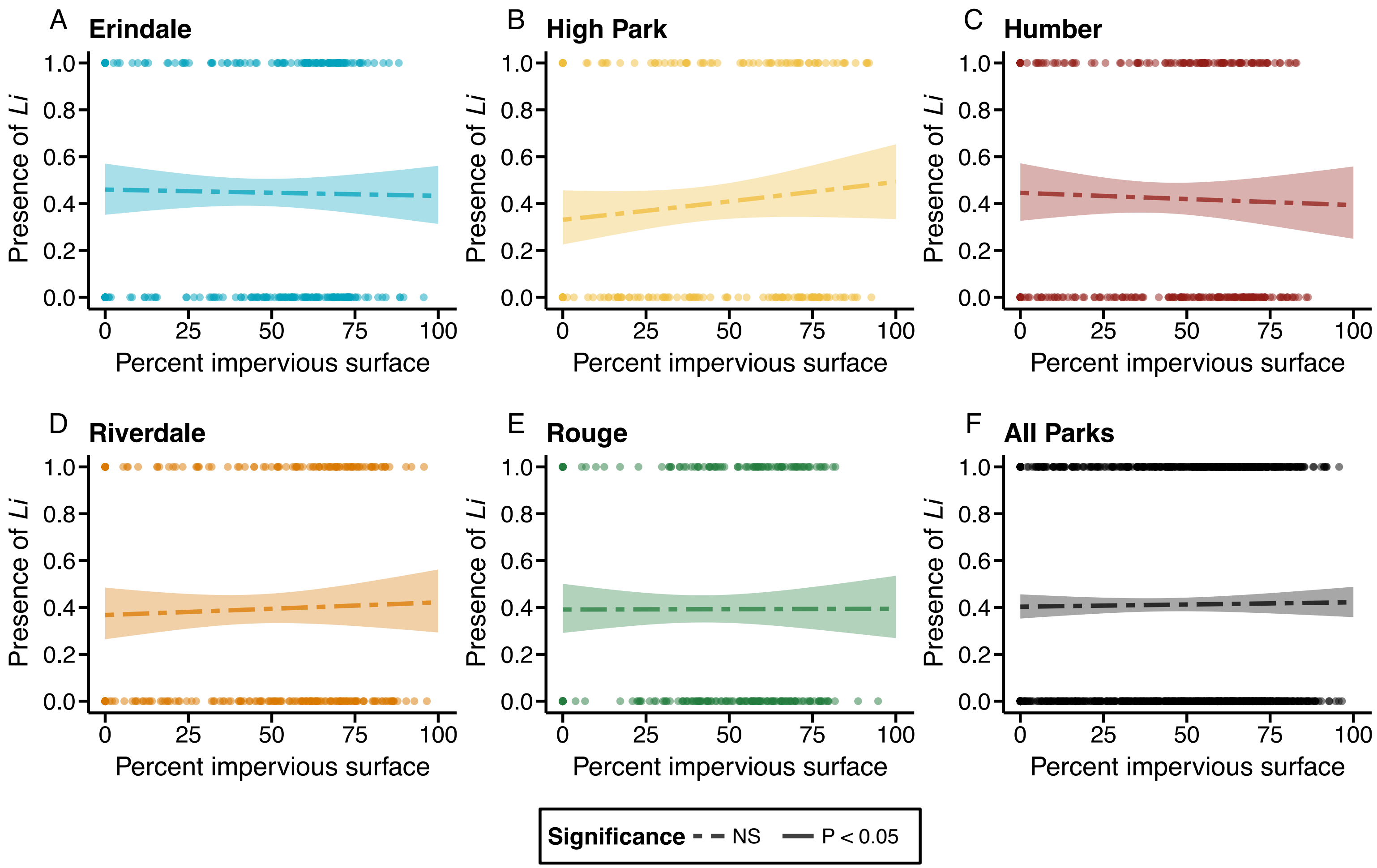


**Figure S3**: Clines in the frequency of *Li* for each of the sampled sites (A – E) and across all sites combined (F). Solid lines represent significant clines ($P < 0.05$) based on a logistic regression of *Li* presence/absence against percent impervious surface. Points represent individual plants. Shaded regions around lines represent the 95% confidence intervals around the fitted models
